# Visual Feedback Weakens the Augmentation of Perceived Stiffness by Artificial Skin Stretch

**DOI:** 10.1101/2020.07.22.215715

**Authors:** Mor Farajian, Hanna Kossowsky, Raz Leib, Ilana Nisky

## Abstract

Tactile stimulation devices are gaining popularity in haptic science and technology – they are lightweight, low-cost, can be easily made wearable, and do not suffer from instability during closed loop interactions with users. Applying tactile stimulation in the form of stretching the skin of the fingerpads, concurrently with kinesthetic force feedback, has been shown to augment the perceived stiffness during interactions with elastic objects. However, all of the studies to date have investigated the perceptual augmentation effects of artificial skin-stretch in the absence of visual feedback. We investigated how visual displacement feedback affects the augmentation of perceived stiffness caused by the skin-stretch. We used a forced-choice paradigm stiffness discrimination task with four different conditions: force feedback, force feedback with artificial skin-stretch, force and visual feedback, and force and visual feedback with artificial skin-stretch. We found that visual displacement feedback weakens the skin-stretch induced perceptual augmentation and improves the stiffness discrimination sensitivity.

## I. Introduction

During everyday interactions with objects, we receive information from multiple senses and integrate them to form our perception of the objects’ mechanical properties, such as the stiffness of the object. The stiffness of an object is defined as the ratio between an applied force on the object and the displacement produced by this force. Since we do not possess stiffness sensors, the perception of stiffness requires high level integration of displacement and force information [1]–[3]. These two components, displacement and force, are sensed by both the visual and haptic senses [4]. The visual information can provide feedback on the amount of displacement through length estimation, and on the force magnitude through surface deformation. The haptic information is the information received through the sense of touch and is comprised of two modalities - kinesthetic and tactile. The kinesthetic information is sensed by muscle spindles (length and shortening velocity of the muscle) and Golgi tendon organs (tension in the tendon). Tactile information is sensed by cutaneous mechanoreceptors that respond to a deformation of the skin, e.g. stretch and vibration [5]. Supplying both visual and haptic information may enhance the accuracy of the position and force estimations relative to what would be achieved using one sense alone. However, while the integration of tactile and kinesthetic information when forming stiffness perception was addressed in several recent sduties [6], [7], the effect of visual feedback on this integration is not known to date.

The nervous system integrates information received from multiple sensory sources to form a single percept. This allows for a robust system that can substitute one sense for another in the event of missing or degraded information, and improves perceptual accuracy compared to what would be possible to achieve with individual sensory sources of information [8]. Previous studies proposed that there are cases in which this integration is in accordance with Maximum Likelihood (MLE). That is, cases where the integration is a convex combination that weights different senses proportionally to their reliability, and is therefore statistically optimal. For example, Ernst and Banks found that haptic and visual information are optimally integrated when forming height perception. They showed that adding varying levels of noise to the visual input increases the uncertainty associated with this modality, thus lowering the weight attributed to it by the nervous system [9]. Furthermore, visual and haptic information have also been shown to be optimally integrated when forming shape perception [10]. In a stiffness magnitude estimation experiment, Varadharajan et al. [11] found that the discrimination accuracy of the participants increased by over 20% when combining haptic and visual information relative to relying on just haptic information. In an additional study, Wu et al. [12] examined the influence of visual feedback on the perceived stiffness of virtual compliant buttons and concluded that the two senses are fused in an optimal manner. They showed that in the absence of visual feedback, participants compliance estimates were biased such that objects that were farther felt softer. Their results indicate that presenting participants with visual information reduces this haptic bias and causes participants’ perceptual judgments to be more reliable.

Contrary to the aforementioned studies, there are cases in which the integration between visual and haptic information has been found not to be optimal. When two sources of information are integrated according to MLE, the relatability of the outcome is increased as measured by a decrease in perceptual variance. Thus, when the variability of the combined estimation is higher than the variability of either one of the individual estimations, this can serve as an indication of sub-optimal integration. For example, Drewing et al. showed that the variability of the stiffness estimation created using both visual and haptic information was similar to the variability of an estimation created using haptic information, suggesting sub-optimal integration between these two senses when forming stiffness perception [13]. This finding is supported by several other studies which showed sub-optimal integration that is biased toward vision when combined with haptic information to form stiffness perception [4], [13], [14].

In all of these studies, when both visual and haptic information were presented, the focus was on kinesthetic force feedback. However, haptic information is comprised of kinesthetic and tactile information, both of which contribute to the perception of stiffness. To study the separate roles of each of the two haptic modalities, technologies which allow for the uncoupled application of kinesthetic and tactile information are required. Toward this end, several different devices for tactile stimulation to the finger-pads have been developed [6], [15], [16]. Using such devices, skin-stretch was used to convey direction [18], to augment friction [15], and to substitute the kinesthetic information in navigation tasks [19]. Moreover, in a sensory substitution study, a tactor-induced skin-stretch device was successfully used to convey stiffness information in a teleoperated palpation task, and it was more accurate than the widely-used vibration feedback [17]. Studies have also shown that adding artificial skin-stretch augments the perceived stiffness, and that this augmentation is a linear function of the amount of stretch [6], [7]. Hence, these devices enable the augmentation of perception without increasing the kinesthetic force, and can therefore be beneficial in the context of providing high levels of force feedback while taking practical limitations, such as closed-loop stability, into account.

Quek et al. [6] hypothesized that the perceptual augmentation in stiffness perception caused by the addition of the artificial skin-stretch may be the result of integrating tactile and kinesthetic information to form the perception of force. However, these studies did not provide participants with visual feedback about object deformation, which has the potential to affect the integration of the haptic information. For example, in [20], the addition of visual feedback influenced the effect of introducing delay into force feedback on the control of grip force in a stiffness discrimination task. The effect of adding visual feedback when both kinesthetic and artificial tactile feedback are presented is currently unknown. Investigating different combinations of visual, kinesthetic and tactile feedback can shed light on how they are combined in the nervous system. Moreover, in many practical scenarios haptic information is presented together with visual information, and therefore, this understanding is imperative for the effective design of haptic feedback devices for applications with physical human-robot interaction, such as teleoperation and robot-assisted surgery [17], [21], [22].

In this work, we studied the influence of visual deformation feedback on the perceptual augmentation of stiffness that is caused by adding artificial skin-stretch to force feedback. Participants probed pairs of virtual force fields that emulated elastic objects, and determined which had a higher level of stiffness. In one of two sessions, participants received visual feedback about the deformation of the probed force field, and in the other, no visual feedback was provided. In both sessions, in half of the trials, artificial tactile feedback was applied during the probing of one of the force fields. We hypothesize that adding visual deformation feedback may enhance participants’ ability to perceive stiffness accurately and precisely. Visual feedback contributes to veridical information about the stiffness of the force field. Therefore, we expect to see a decrease in the overestimation of the perceived stiffness caused by the artificial skin-stretch when participants receive visual feedback. In addition, we expect that visual feedback may enhance their ability to distinguish between smaller differences between stiffness levels, and thus, decrease participants’ uncertainty relative to conditions without visual feedback.

## Methods

### A. Participants

21 participants right handed participants (14 females, ages 21-29) conducted the experiment after signing an informed consent form approved by the Human Subject Research Committee of Ben-Gurion University (BGU) of the Negev, Be’er Sheva, Israel. The participants were compensated for their participation, regardless of their success or completion of the experiment.

### B. Experimental Setup

The participants sat in front of a virtual reality system and used the thumb and index finger of their right hand to grasp a skin-stretch device that was mounted on a PHANTOM® Premium 1.5 haptic device (3D SYSTEMS) [Fig. 1]. Two tactors came into contact with the skin of the fingers and moved in the vertical direction to stretch the skin through tactor displacement [Fig. 1(c)]. The skin-stretch device was identical to the one that is described in detail in [7]. The participants looked at a semi-silvered mirror showing the projection of an LCD screen placed horizontally above it. An opaque screen was fixed under the mirror to block the participants’ view of their hands. During the experiment, the participants wore noise cancelling headphones (Bose QC35) to eliminate auditory cues.

**Fig. 1.**
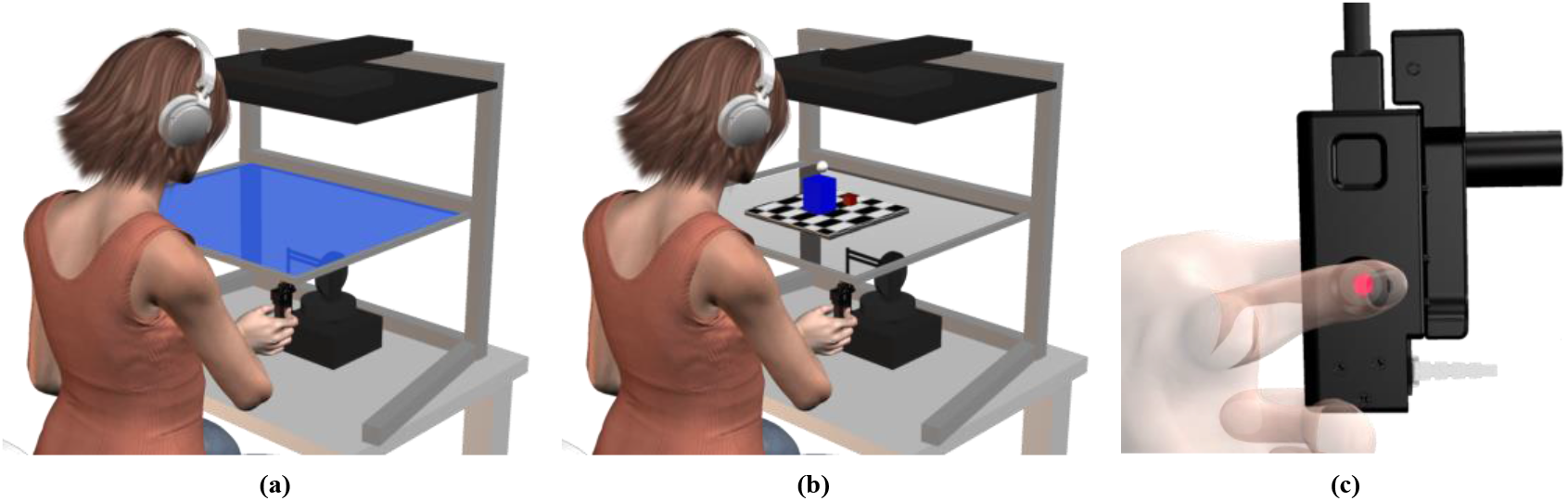
Experimental setup: the participants sat in front of a virtual reality rig, and held the skin-stretch device, which was mounted on the end of a haptic device. (a) The screen participants saw during conditions without visual feedback: the force field was indicated to the participants by the screen color, which was either red or blue. (b) The screen participants saw during conditions with visual feedback: the force field was displayed as a cube sitting on a checkerboard-patterned floor, and the color of the cube was either red or blue. In both conditions, (a) and (b), the screen was completely opaque, but is presented as semi-transparent in this figure to visualize the haptic device beneath it. (c) Side view of the skin-stretch device. The participants used the thumb and index finger of their right hand to grasp the device. Two tactors (red rod) came into contact with the skin of the fingers and moved in the vertical direction to stretch the skin through tactor displacement.

The haptic device applied kinesthetic force feedback with natural skin-stretch, and the skin-stretch device created the additional artificial tactile stimulation. The force and stretch were proportional to the vertical position of the end of the haptic device and were applied only when participants were in contact with the force field, which was defined to be the negative half of the vertical axis:

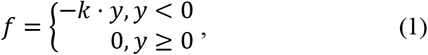

where k [N/m] is the stiffness, and y [m] is the penetration distance into the virtual force field. We used the skin-stretch device to apply tactile stimuli by means of tactor displacement:

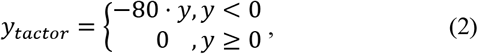

where 80 [mm/m] is the tactor displacement gain, and y [m] is the same penetration distance into the force field.

### C. Protocol

In a forced-choice stiffness discrimination task, participants probed pairs of virtual elastic force fields, designated *standard* and *comparison*, and chose which force field had a higher level of stiffness. Participants could probe the force fields as many times as they desired and switch freely between them before making their decision. In some trials, visual feedback about the force field deformation was presented to the participants. Additionally, there were trials in which artificial tactile stimulation was applied. This led to four experimental conditions which differed in the visual feedback that the participants received and the presence of tactile stimulation: (1) force feedback (F), (2) force feedback with artificial skin-stretch (FS), (3) force and visual feedback (FV), and (4) force and visual feedback with artificial skin-stretch (FVS).

The *standard* force field had a constant stiffness value of 85 N/m and the skin-stretch gain was 80 mm/m in conditions with skin-stretch stimulation. This skin-stretch gain was chosen based on our previous study [7], which demonstrated a clear augmentation in the perceived stiffness due to this level of stretch. The stiffness level of the *comparison* force field was selected from a range of 10 values, evenly spaced between 40-130 N/m. During trials without visual feedback, each force field (*standard* and *comparison*) was indicated to the participants by the screen color, which was either red or blue [Fig. 1(a)]. In these trials, participants did not receive any feedback on the amount of penetration distance into each of the force fields. To switch between the two force fields, participants were instructed to raise the end of the robotic device to at least 30 mm beyond the boundary of the force field. During trials with visual feedback, the force field was displayed as a large cube sitting on a checkerboard-patterned floor, and the color of the cube was either red or blue. The position of the hand was displayed as a white sphere which participants used to press down on the force field. As a result, the force field deformed according to participant’s penetration distance and the stiffness level of the force field. To switch between the two force fields, participants placed the white sphere on a virtual brown button, represented on the screen as a little cube that was placed to the side of the force field [Fig 1(b)]. The visual feedback was veridical with the hand movement and was presented in both the *standard* and *comparison* force fields. In both cases (trials with and without visual feedback), after the participants interacted with the force fields and decided which one was stiffer, they pressed a keyboard key that corresponded to the screen or cube color of the stiffer force field.

In total, there were 10 *comparison* stiffness levels and four *standard* conditions, amounting to a total of 40 *standard*-*comparison* pairs, each of which was repeated eight times throughout the experiment. Each participant therefore performed 360 trials: 40 training trials (to allow participants to become familiarized with the experimental setup), and 320 test trials. The experiment was divided into two sessions that were completed over two days of 180 trials each; half the participants (Group 1, N=11) completed conditions F and FS on the first day and conditions FV and FVS on the second day, while the other half (Group 2, N=10) completed the two sessions in the opposite order.

### D. Data Analysis

For each of the 21 participants, we used the Psignifit toolbox 2.5.6 [23] to fit psychometric curves to the probability of responding that the *comparison* force field was stiffer as a function of the difference between the stiffness levels of the *comparison* and the *standard* force fields. We repeated this procedure for the four experimental conditions, and computed the point of subjective equality (PSE) and the just noticeable difference (JND) of each psychometric curve. The PSE is defined to be the stiffness level at which the probability of responding that the *comparison* force field had a higher level of stiffness was 0.5:

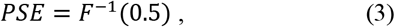

where *F* is the psychometric function, a logistic function in our case. A positive PSE value (that is, a rightward shift of the psychometric curve) represents an overestimation of the *standard* force field stiffness, and a negative PSE value (a leftward shift) indicates underestimation. The JND is measured as half of the difference between the *comparison* stiffness levels corresponding to the 0.75 and the 0.25 probabilities of choosing that the *comparison* force field was stiffer:

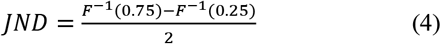

The JND quantifies the sensitivity of the participant to small differences between the stiffness levels of the two force fields, and therefore is an indication of the uncertainty experienced by the participants when choosing which force field was stiffer. The PSE value in the FS condition of one of the participants from Group 1 was close to 100 N/m. This increase exceeded the range of tested *comparison* stiffness levels, and based on our work in [7], is considered an outlier, and therefore, this participant was excluded from the analyses. Nevertheless, this outlier removal did not qualitatively change any of our conclusions.

After computing the PSE and JND values of each of the participants in each of the conditions, we examined the effect of the different conditions on these two values using a repeated-measures General Linear Model. The dependent variables in the two separate analyses were the PSE and JND values, and the independent variables were the stretch stimulus (categorical, df=1), the visual feedback (categorical, df=1), and the participants (random, df=19). The statistical model also included the interaction between the stretch stimulus and the visual feedback. To assess if there was a difference between the two groups, we used a nested General Linear Model, where the participants variable was nested in the group-number variable (categorical, df=1). The main interest of this study was the interaction between stretch stimulus and visual feedback, which if significant would lead to performing three t-tests to compare between the different conditions (F and FS, FV and FVS, and FS and FVS) with the Holm-Bonferroni method to correct for errors stemming from multiple comparisons. We present the p-values after this correction, and therefore the threshold significance level in all the reported tests is 0.05.

## II. Results

The psychometric curves of a typical participant are shown in Fig. 2(a). The orange curves represent conditions without visual feedback, and the blue curves represent conditions with visual feedback. The light colors represent conditions without tactile stimulation and the dark colors represent conditions with tactile stimulation. As shown in the psychometric curves, in trials without tactile stimulation (light blue and orange curves), the PSE was close to zero, and the slope of the psychometric curve (as quantified by the JND) was steep, indicating that the participant could accurately distinguish between the stiffness levels of the two force fields. Stretching the finger-pad skin (dark blue and orange curves) led to a rightward shift of the psychometric curves, and a positive PSE, indicating an overestimation of the perceived stiffness. Although this perceptual augmentation effect was observed in both the FS and FVS conditions, the presence of the visual feedback decreased this effect. That is, the increase in the perceived stiffness in the FVS condition was smaller than in the FS condition (a smaller rightward shift of the dark blue curve relative to the dark orange curve).

**Fig. 2.**
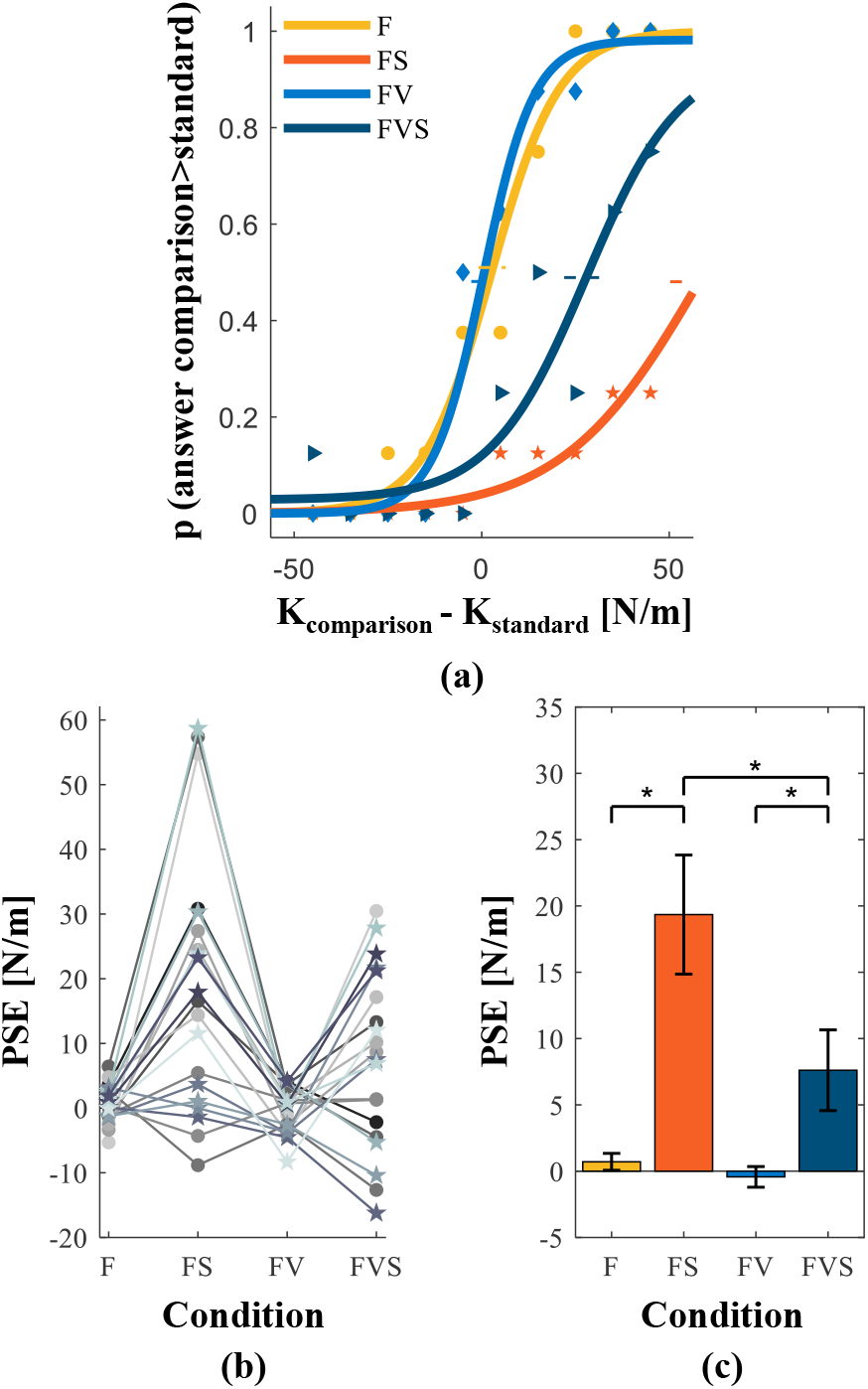
(a) An example of the psychometric curves of a typical participant in the different conditions (F,FS,FV,FSV). The abscissa is the difference between the stiffness levels of the *comparison* and the *standard* force fields, and the ordinate is the probability of responding that the *comparison* force field had a higher level of stiffness. The horizontal dashed lines represent the standard errors for the PSE values. (b) The PSE values of all the participants as a function of the different conditions. The circles and lines colored in different shades of gray represent the data of Group 1 (N=10), and the stars and lines colored in different shades of blue represent the data of Group 2 (N=10). (c) The averaged PSE values across all the participants (N=20), as a function of the different conditions. The black bars represent the standard errors, and the asterisks indicate a statistically significant difference (p<0.05).

The PSE values of all the participants in the different conditions are shown in Fig. 2(b-c). The results of all the participants were similar to those of the typical participant; when participants were exposed to artificial skin-stretch, the PSE increased relative to the conditions without the stretch stimulus (main effect of ‘stretch stimulus’: *F*_(1,57)_ = 27.90, *p* < 0.0001). The addition of visual feedback led to a decrease in the PSE values relative to the PSE in the conditions without the visual feedback (main effect of ‘visual feedback’: *F*_(1,57)_ = 6.50, *p* = 0.0135). In addition, the interaction between the stretch stimulus and the visual feedback variables was also statistically significant (interaction between ‘stretch stimulus’ and ‘visual feedback’: *F*_(1,57)_ = 4.40, *p* = 0.0403). That is, when the visual deformation feedback was not presented, skin-stretch augmented the perceived stiffness [6], [7], but the presence of visual feedback decreased this perceptual augmentation (*t*_FS−FVS_ = 3.28, *p*_FS−FVS_ = 0.0017). Additionally, the other two planned t-tests we conducted revealed significant differences between the PSE values in the F and FS conditions (*t*_F−FS_ = 5.21, *p*_F−FS_ < 0.0001) and the FV and FVS conditions (*t*_FV−FVS_ = 2.25, *p*_FV−FVS_ = 0.0283).

The JND values of each of the participants, and their means, in the four different conditions are depicted in Fig. 3(a) and 3(b) respectively. The stretch stimulus led to an increase in the JND values relative to the JND in the conditions without the stretch stimulus (main effect of ‘stretch stimulus’: *F*_(1,57)_ = 5.44, *p* = 0.0232). That is, the stretch stimulation may have increased uncertainty about the cutaneous information, and therefore increased the uncertainty experienced by the participants when choosing which force field was stiffer. When comparing between the FS and FVS conditions, we found that participants were more accurate in distinguishing between the stiffness of two force fields in the FVS condition (as indicated by a lower JND). However, this effect was not statistically significant (interaction between ‘stretch stimulus’ and ‘visual feedback’: *F*_(1,57)_ = 0.28, *p* = 0.6014).

**Fig. 3.**
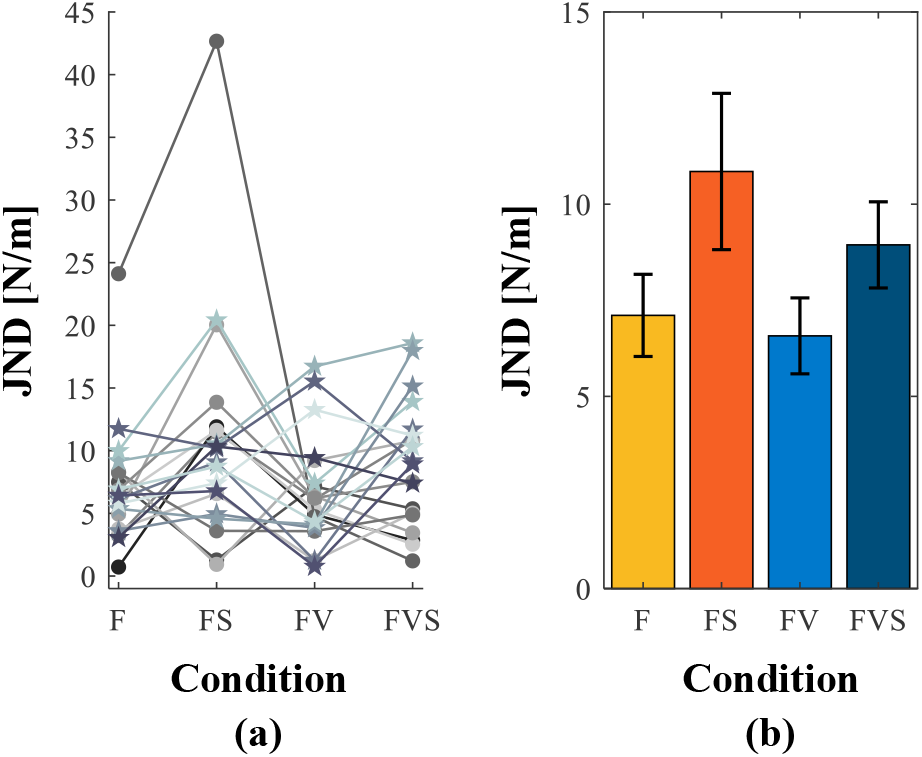
(a) The JND values of all the participants as a function of the different conditions (F,FS,FV,FVS). The circles and lines colored in different shades of gray represent the data of Group 1 (N=10), and the stars and lines colored in different shades of blue represent the data of Group 2 (N=10). (b) The averaged JND values across all the participants (N=20), as a function of the different conditions. The black bars represent the standard errors.

This experiment was divided into two sessions; Group 1 completed conditions F and FS on the first session and condition FV and FVS on the second session, while Group 2 completed the two sessions in the opposite order. To test if there was an effect of the order of sessions on the results, we compared between the results of the two groups and found no significant differences between them (rm-General Linear Model, PSE: main effect of ‘group number’: *F*_(1,18)_ = 0.07, *p* = 0.8015; JND: main effect of ‘group number’: *F*_(1,18)_ = 0.79, *p* = 0.3852). Therefore, and because the order was balanced, we combined the results of the two groups in all the statistical analyses in the paper.

## III. DISCUSSION

In this study, we examined the effect of visual feedback on the perceptual augmentation caused by adding artificial skin-stretch to force feedback. Our results show that the addition of the visual information reduces the skin-stretch augmentation, indicating that visual feedback improves the ability of the participants to accurately perceive stiffness. Additionally, we observed a decrease in the JND in conditions with both haptic and visual feedback relative to those with haptic feedback alone. While this decrease was not statistically significant and requires further investigation, it hints that the addition of visual feedback to skin-stretch stimulus during interactions with elastic force fields may enhance the discrimination accuracy.

Previous studies demonstrated that when both haptic and visual information are presented, weight is attributed to the information received from each of the senses to form a single percept [8], [24]. Additionally, in the event of a discrepancy between the visual and haptic information, the estimate of the perceived stiffness was biased toward the visual information [4], [13], [14]. In this work, participants received either haptic feedback alone, or both visual and haptic feedback. The haptic feedback was comprised of kinesthetic force and artificial tactile skin-stretch, which has been shown to augment the perceived stiffness [6], [7]. We showed that presenting participants with visual feedback, which was consistent with the force feedback, lowered this perceptual augmentation effect, indicating that less weight was attributed to the haptic information. A comparison between two different works [6], [7] that studied the increase in the perceived stiffness caused by artificial skin-stretch supports these findings. Both studies reported an augmentation in the perceived stiffness, however in [6] the inter-subject variability was larger. A potential explanation is that in [7] participants received no visual feedback, whereas in [6] partial visual feedback was provided, supporting our claim that visual feedback may affect the perceptual illusion. Wu et al. [12] found that adding visual displacement information to haptic information reduced a haptic bias to feel more distant objects as softer. Varadharajan et al [11] found no significant contribution of vision to the perception of stiffness magnitude during interaction with haptic-visual conditions. To conclude, many studies have demonstrated the effect of visual feedback on stiffness perception, however, the contribution of visual information to the formation of a single percept might be task depended.

In addition to the effect of visual information on the perceived stiffness, the addition of this sensory input has also been shown to affect the variance of the stiffness estimate. Visual information has been shown to be a more reliable source of information than haptic feedback, and when presented together with haptic feedback, lowered the variance of the perceptual estimates, leading to smaller JND values [4], [9], [24]. Kuschel et al. [3] presented participants with both haptic and visual compliance information and found that when the visual and haptic inputs were congruent, the perceived compliance was a weighted combination of the two sensory inputs such that the variance of the final estimate was minimized (i.e., statistically optimal). However, when the visual input was distorted relative to the haptic information, the integration was not optimal. Additionally, Varadharajan et al [9] concluded that adding a visual rendering of the interaction in a stiffness discrimination task improved the discrimination accuracy. Our results correspond to those of these studies; the discrimination accuracy of the participants during trials with force and artificial skin-stretch with visual feedback was lower than in trials with force feedback with artificial skin-stretch and no visual feedback. However, the effects in our study were small and not statistically significant.

Another interesting observation is the decrease in the discrimination accuracy in the conditions with the tactile stimulus relative to the conditions without this stimulus. This result is consistent with our previous work [7] in which we studied the effect of applying different levels of skin-stretch on the perceived stiffness. We found that increasing the magnitude of stretch decreased the discrimination accuracy of some of the participants. A possible explanation for this result is that the artificial skin-stretch may have introduced uncertainty into the haptic information and decreased participants' sensitivity to small differences between the stiffness levels of the two force fields. However, the way in which participants interpret this tactile stimulus, as additional stretch or maybe as noise that leads to increased stimulation of the mechanoreceptors, is an open question.

The initial goal of this work was to study the contribution of visual deformation feedback to the perceptual augmentation caused by adding artificial skin-stretch to force feedback. However, this work can also shed light on the integration between the visual feedback and the two haptic modalities. According to optimal integration, when discrepant haptic and visual information are presented, the perceived stiffness would be a weighted average of the estimates created using each of the sensory inputs. Additionally, the discrimination accuracy in the haptic-visual condition would be higher than that in the haptic alone condition [4], [9], [24]. The results of our work are consistent with these predictions; participants maybe interpret the artificial skin-stretch as a noisy source of information and therefore lowered the weight attributed to the haptic information. As a result, relatively more weight may have been given to the visual source of information. In addition, consistently with optimal integration, the addition of the visual feedback improved the discrimination accuracy of the participants, albeit surprisingly, this effect did not reach statistical significance. Although our results are compatible with optimal integration, our experiment does not allow for the calculation of the weights attributed to each of the senses and therefore further investigation with more experimental conditions is needed to confirm this.

Understanding the integration between visual, kinesthetic and tactile information can lead to improvements in human-robot interaction applications that currently do not present haptic feedback to the user. Even though the literature is ambiguous about how the performance of the user benefits from receiving haptic feedback in cases where visual feedback is presented [25], [26], there are likely more than a few tasks that cannot be done without both sensory inputs. Therefore, designing systems that provide users with sensory feedback that is tailored to their natural information processing strategies can benefit from calculated combination of visual feedback and haptic feedback comprised of kinesthetic and tactile information.

## Acknowledgment

We thank Nitsan Tirza for assistance in data collection.

